# New High Throughput Approaches to Detect Partial-body and Neutron Exposures on an Individual Basis

**DOI:** 10.1101/646711

**Authors:** Igor Shuryak, Helen C. Turner, Jay R. Perrier, Lydia Cunha, Monica Pujol Canadell, Mohammad H. Durrani, Andrew Harken, Antonella Bertucci, Maria Taveras, Guy Garty, David J. Brenner

**Affiliations:** Center for Radiological Research, Columbia University Irving Medical Center, New York, NY, USA

## Abstract

Biodosimetry-based discrimination between homogeneous total-body photon exposure and complex irradiation scenarios (partial-body shielding and/or neutron + photon mixtures) can improve treatment decisions after mass-casualty radiation-related incidents. Our study objective was to use high-throughput biomarkers to: a) detect partial-body and/or neutron exposure on an individual basis, and b) estimate separately the photon and neutron doses in a mixed exposure. We developed a novel approach, where metrics related to the *shapes* of micronuclei distributions per binucleated cell in *ex-vivo* irradiated human lymphocytes (variance/mean, kurtosis, skewness, *etc*.) served as predictors in machine learning or parametric analyses of the following scenarios: (A) Homogeneous gamma-irradiation, mimicking total-body exposures, *vs.* mixtures of irradiated blood with unirradiated blood, mimicking partial-body exposures. (B) X rays *vs.* various neutron + photon mixtures. Classification of samples as homogeneously *vs.* heterogeneously irradiated (scenario A) achieved a receiver operating characteristic curve area (AUROC) of 0.931 (uncertainty range of 0.903-0.951), and R^2^ for actual *vs.* reconstructed mean dose was 0.87. Detection of samples with ≥10% neutron contribution (scenario B) achieved AUROC of 0.916 (0.893-0.943), and R^2^ for reconstructing photon-equivalent dose was 0.77. These encouraging findings demonstrate a proof-of-principle for the proposed approach of analyzing micronuclei/cell distributions to detect clinically-relevant complex radiation exposure scenarios.

## Introduction

The need for high-throughput biodosimetry in response to a large-scale radiological event such as improvised nuclear device (IND) detonations stems from several considerations ^1^. First, triage, which is expected to take place away from a hospital, is crucial for preventing treatment locations from being overwhelmed. Second, to determine the optimal treatment for an individual exposed to a large radiation dose, it is critical to quantitatively reconstruct the radiation dose that the individual received to identify among an exposed population those individuals who are most likely to develop acute or late radiation injury. Third, the need to convey credible information about radiation doses to individuals as quickly as possible is a major lesson learned from earlier radiological events ^2^.

It is likely that large numbers of victims of an IND will receive partial body exposure, due to shielding by buildings or vehicles, as well as a mixture of densely ionizing neutrons and sparsely ionizing gamma rays, with the radiation quality and type of exposure varying between individuals ^3^. A rapid assessment of exposure type would need to be made for a large number of individuals following a large-scale radiological event. Consequently, there is a well-recognized need for development and utilization of high-throughput assays that can discriminate between these complex irradiation scenarios like partial-body shielding and/or neutron + photon mixtures from simpler exposures such as homogeneous total-body photon exposure ^4–6^. Such reconstruction of exposure type is important for making appropriate triage and treatment decisions in mass casualty situations.

### Significance of neutrons

A likely scenario for an IND is a gun-type detonation using highly enriched uranium ^7^. Here, the prompt exposure will consist of gamma rays combined with a device-dependent dose of fast neutrons ^8^. Monte-Carlo based estimates of the neutron component from a 10 kT urban ground burst IND ^3^ suggest in-air neutron fractions of 20% to 90% and, more relevantly, organ-dose neutron fractions of 3 to 14% in the colon and 6 to 27% in the bone marrow ^3^. Due to the high relative biological effectiveness of neutrons for causing cytogenetic damage ^9–11^, the neutron dose contributes roughly 4 times the damage of an equivalent photon dose. Consequently, these neutron components are likely to have a profound impact on disease type and progression ^12–15^. It is also likely that different countermeasures will be required for neutron-induced disease and photon-induced exposure ^16^

### Significance of partial body exposures

A significant proportion of individuals exposed indoors to the initial blast from an IND will be exposed non-homogeneously, to a partial body exposure, due to shielding by objects and building materials ^7^. By contrast, external fallout is likely to result in a more homogenous exposure but decreases over time, approximately following a power function called “the 7:10 rule” ^17, 18^. Partial body exposure has important consequences in terms of medical countermeasures and disease progression. For example, the hematopoietic system can recover much better after high-dose irradiation when part of the body containing bone marrow (*e.g.* one or more limbs) is shielded ^19^. In animal studies, even 5% bone marrow shielding results in a large increase in survival from hematopoietic acute radiation syndrome (H-ARS) ^20^ and can also profoundly affect the GI syndrome ^21^. A simple biodosimetric dose reconstruction that estimates a single dose number assumes uniform irradiation and would thus generate incorrect results, likely overestimating the risk for H-ARS and underestimating the risk for later disease in the organs that were irradiated.

### Current approaches for evaluating complex exposures

There is a large body of literature on various computational biodosimetry approaches for estimating radiation doses in various exposure scenarios based on micronuclei yields and other cytogenetics markers like dicentric chromosomes ^22–32^. As compared to uniform photon exposures, both neutron and partial body exposures result in a non-poissonian distribution of damage in the cells scored for biodosimetry due to the shielding and/or differences in radiation track structure and energy deposition patterns. Thus, there are a larger number of undamaged cells than would be expected based on Poisson statistics, coupled with more damage in those that are traversed by ionizing tracks. While these phenomena has been observed for many years and applied to the dicentric assay, essentially by analyzing the proportion of undamaged metaphases ^33–35^, these approaches can be directly applied to high-throughput assays ^5, 6^.

It is well known that cytogenetic damage distribution shapes, which are commonly modeled by Poisson, Negative Binomial or Neyman distributions ^26, 36, 37^, can change depending on exposure type. For example, densely ionizing radiations like neutrons tend to produce “overdispersed” distributions of cytogenetic damage, where the ratio of variance/mean becomes significantly higher than in a standard Poisson distribution ^38, 39^. Partial-body exposures also tend to produce overdispersion because even if the damage distribution for a homogeneous exposure is Poisson, the contribution from a shielded fraction of the body that received a much lower dose would cause the distribution to become a mixture of two or more Poissons with different means ^40, 41^. Although the methodologies for analyzing these phenomena differ (*e.g.* frequentist *vs.* Bayesian techniques), a more common popular approach is to fit selected probability density functions (*e.g.* Zero-Inflated Poisson or Negative Binomial) to the data ^31, 42, 43^. The best-fit parameters and their uncertainties are then used to estimate the outcomes of interest. However all these techniques are based on manual scoring of the number of dicentrics in a large number of metaphases and they are not compatible with processing of tens of thousands of samples using automated biodosimetry.

### Potential of machine learning techniques

The data fitting methodologies described above rely on parametric regression, such as linear or linear quadratic functions, to describe the radiation response. To our knowledge, ensemble machine learning techniques such as random forests (RF) and generalized boosted regression models (GBM) ^44, 45^ have not been used for radiation biodosimetry applications. Both parametric and machine learning regression approaches have specific advantages and disadvantages. Parametric models are easily interpretable because each fitted coefficient has a specific meaning for relating a given predictor or predictor combination to the outcome(s). RF and GBM can be more complicated to interpret because they consist of multiple (usually >100) decision trees. However, RF and GBM tend to be more flexible than parametric models in describing nonlinear dependences and interactions between predictors, and therefore tend to be more accurate.

Ensemble methods like RF and GBM train and test multiple models of a given type on randomly-selected subsets of the analyzed data set and combine the results, thereby generating more robust and accurate predictions than those obtainable using a single model ^45^. RF uses decision trees as base models, and employs “bagging” and tree de-correlation approaches to improve performance. The bagging (bootstrapping and aggregation) procedure involves generating bootstrapped samples and using a random subsample of the features for each fitted decision tree. Decision trees have some very useful properties for analyzing data set types such as those in the current application. For example, they are not sensitive to outliers and to the presence of many weak or irrelevant predictors. They are also unaffected by monotonic (*e.g.* logarithmic) transformations of the data. RF readily allows for multivariate analysis with more than one outcome variable and a common set of predictor variables. All of these properties can potentially prove useful in biodosimetry applications. GBM also uses decision trees, but the trees are averaged by boosting rather than bagging. Boosting involves iterative fitting of trees: the data are reweighted so that the next trees focus more strongly on those data points on which previous trees performed the worst. GBM readily accommodates different types of error distributions, e.g. Gaussian for continuous data and Bernoulli for binary data.

### Study design

In this work, we employed machine learning approaches in a novel role, using the *shape* of the distribution curve of micronuclei per binucleated cell as a source of information for discriminating between simple and complex radiation exposure scenarios, *e.g.* total-body *vs.* partial-body photon exposures, or *vs.* neutron + photon mixtures. Specifically, using our high-throughput CBMN assay ^5, 9, 46^ we wish to evaluate on an individual basis: 1) the photon and the neutron doses and the fraction of neutrons in the total dose after a mixed exposure, 2) whether there was indeed a partial body exposure. Our study design (shown schematically in Fig. 1) consisted of using fresh human peripheral blood samples irradiated ex vivo to analyze the following simple and complex exposure scenarios:

**Figure 1.**
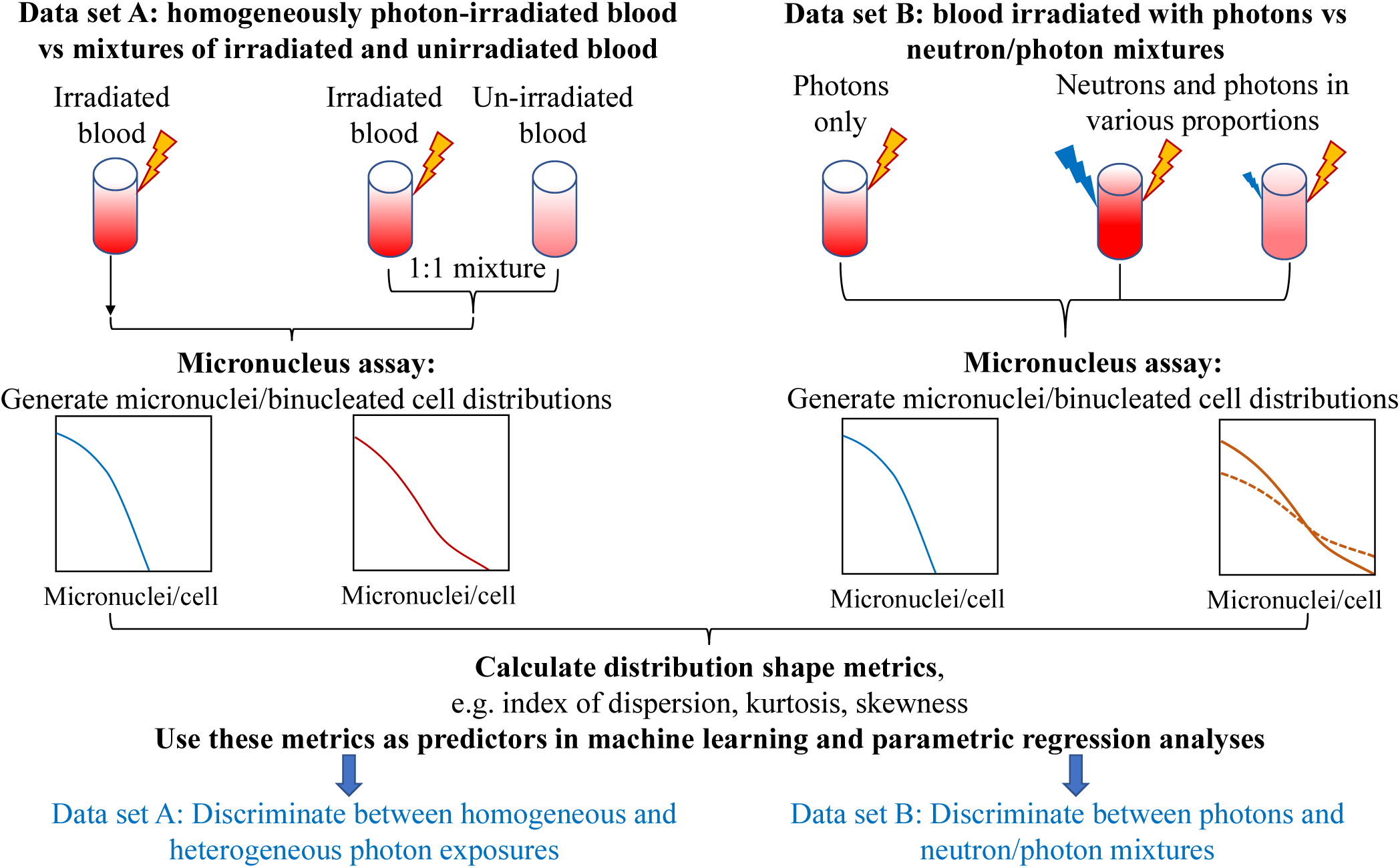
A schematic representation of our study design aimed at developing new computational methods for discriminating between triage-relevant simple and complex radiation exposure scenarios. We used ex vivo irradiated human blood to generate two data sets (A and B), and analyzed each of them using a novel application of machine learning techniques. The data sets and analysis methods are described in detail in the Materials and Methods section. Yellow lightning symbols indicate photon irradiation, and blue ones indicate neutron irradiation of blood samples. Curves of various colors indicate probability distributions of micronuclei per cell, where the y-axis is probability density. Solid *vs.* dashed lines indicate the effects of different neutron proportions. These schematic distributions are intended to illustrate that complex exposure scenarios, such as mixtures of irradiated and unirradiated blood, or photon + neutron exposures, produce larger “tails” (*i.e.* larger probabilities of multiple micronuclei per cell) than simple exposures.

*Scenario A.* Homogeneous 0, 2, 4, or 8 Gy gamma irradiation, mimicking total-body exposures, *vs.* 1:1 mixtures of 4 or 8 Gy irradiated blood with unirradiated blood, mimicking partial-body exposures. In this data set, 4 Gy-irradiated blood mixed with unirradiated blood was intended to produce a similar mean micronuclei yield to blood irradiated with a homogeneous dose of 2 Gy. The goal of the computational biodosimetry approach in this scenario was to correctly classify such situations as homogeneous exposures *vs.* mixtures.

*Scenario B.* Photons (0-4 Gy of x-rays) *vs.* mixtures of neutrons + photons in various proportions (up to 3 Gy neutrons). The neutron proportions were intentionally varied over a wide range to mimic various types of realistic exposure scenarios. The goal of the computational biodosimetry approach in this scenario was to distinguish neutron + photon mixtures from pure photon exposures, and to quantify the neutron contribution.

## Materials and Methods

### Blood collection and irradiation

Fresh peripheral blood samples were collected by venipuncture into 6 ml lithium-heparinized Vacutainer^®^ tubes (BD Vacutainer™, Franklin Lakes, NJ) from healthy female and male donors with informed consent as approved by the Columbia University Medical Center Institutional Review Board (IRB protocol no: AAAE-2671). Healthy blood donor volunteers, aged between the ages of 24 and 48 years were non-smokers and in relatively good health at the time of donation with no known exposure to x rays or CT scan within the last 12 months.

#### i) Neutron and x-ray irradiations

These irradiations were performed at the Columbia IND Neutron Facility (CINF) ^9, 47, 48^. Our broad-energy neutron irradiator has been designed to expose blood or small animals to neutron fields mimicking those from an IND. This spectrum, dominated by neutron energies between 0.2 and 9 MeV that mimics the Hiroshima gun-type energy spectrum at a relevant distance (1-1.5 km) from ground zero ^8, 9^, is significantly different from a standard reactor fission spectrum, because the bomb spectrum changes as the neutrons are transported through air. Blood aliquots (1 ml) in 1.4 ml Matrix 2D-barcoded storage tubes (Thermo Fisher Scientific, Waltham, MA) were prepared and either sham-irradiated or exposed to neutrons and x rays at the Radiological Research Accelerator Facility (RARAF). Details of the IND-spectrum neutron irradiator and dosimetry have been described previously ^9, 48^. Briefly, the aliquoted blood samples were placed in adjacent positions on an eighteen position Ferris wheel. The wheel rotates during irradiations and maintains the sample locations at a distance of 17.5cm and an angle of 60 from the beam’s impingement on a thick beryllium target. Neutron irradiations were performed over several runs with 15-30 μA mixed beams of protons and deuterons on the target generating a neutron dose rate of 1.3-2.6 Gy/h with a 18% concomitant dose of gamma rays. To ensure a uniform scatter dose, equivalent tubes containing water were placed in any empty positions on the wheel. Dosimetry for CINF was performed, on the day of the experiment, as described previously ^9^.

For the mixed photon + neutron exposure studies, some blood samples were exposed to x rays following neutron irradiation. This was done using a Westinghouse Coronado orthovoltage x-ray irradiator running at 250-kVp and 15 mA with a 0.5 mm Cu + 1 mm Al filter (Half Value Layer 2 mm Cu). X rays were delivered at a dose rate of 1.23 Gy/min. All tested combinations of x rays and neutrons are shown in the Supplementary_data_file1 online.

#### ii) Gamma ray irradiations

Irradiations for partial body exposures were performed at the Center for Radiological Research, Columbia University Irving Medical Center, New York. Blood aliquots (6 ml) in 15-ml conical bottom tubes (Santa Cruz Biotechnology^®^ Inc., Dallas, TX) were prepared and transported to a Gammacell 40 ^137^Cesium (^137^Cs) irradiator (Atomic Energy of Canada Ltd.). The blood samples were placed in a custom-built 15 ml tube holder and exposed to 0 (control), 2.0, 4.0, or 8 Gy of γ rays at a dose rate of 0.73 Gy/min. The ^137^Cs irradiator is calibrated annually with TLDs and homogeneity of exposure across the sample volume was verified using EBT3 Gafchromic™ film with less than 2% variation within the sample (Ashland Advanced Materials, Bridgewater, NJ). For the heterogeneous exposures, the blood samples were mixed 1:1 (0 Gy and 4 or 8 Gy).

### Micronucleus assay

Whole blood samples from each dose point were cultured in PB-MAX™ Karyotyping media (Life Technologies, Grand Island, NY), and incubated at 37°C, 5% CO_2_, 98% humidity. After 44 h, the media was refreshed with PB-MAX™ media supplemented with cytochalasin B (Sigma-Aldrich LLC, St. Louis, MO) at a final concentration of 6 μg/mL to block cytokinesis. After a total incubation period of 72 h, the cells were harvested. The cells were treated with 0.075 M KCl solution (Sigma-Aldrich, St. Louis, MO) at room temperature for 10 min. After hypotonic treatment, the cells were fixed with fixative (4:1 methanol:glacial acetic acid). The fixed cell samples were stored at 4°C (at least overnight), dropped on slides, allowed to air dry for 10 min and then stained with Vectashield^®^ mounting media containing DAPI (Vector Laboratories, Burlingame, CA). The slides were left overnight at 4°C prior to imaging.

### Imaging analysis and micronuclei scoring

Slides were imaged using a Zeiss fluorescence microscope (Axioplan 2; Carl Zeiss MicroImaging Inc., Thornwood, NY) with a motorized stage and Zeiss 10× air objective. Quantification of micronuclei yields was performed by automatic scanning and analysis with the Metafer MNScore software (MetaSystems, Althaussen, Germany) using the Metafer classifier described in our earlier work ^49^. Images were captured using a high-resolution, monochrome megapixel charge coupled device (CCD) camera. For each sample, more than 1000 binucleated cells were scored and the micronuclei distribution per cell recorded. The values reported by the Metafer MnScore software were the micronuclei counts per binucleated cell, ranging from 0 to 5. The counts in the bin labeled 5 actually represent the sum of counts with values ≥5, as outputted by the Metafer software. These counts were typically low (median = 0, maximum = 16, whereas the median sum of all counts per sample was 461) and the lack of detailed bin information for bins >5 was unlikely to modify the results substantially.

### Compilation of the Data sets

The experimental data analyzed by this study were compiled into two data sets, labeled A and B, which are presented in the Supplementary_data_file1 online. Data set A consisted of a single experimental design with homogeneous 0, 2, 4, or 8 Gy gamma irradiation, mimicking total-body exposures, *vs.* 1:1 mixtures of 4 or 8 Gy irradiated blood with unirradiated blood, mimicking partial-body exposures. Data set B was a large compilation blood samples which had been exposed to IND-spectrum neutrons and neutron + photon mixtures in various proportions (up to ∼82% neutrons), including one previously published sample set ^9^. The goal of combining such a large number of experiments was to increase statistical power and to clarify the main patterns of interest, such as the dependences of micronuclei per cell distributions on photon and neutron contributions in the dose.

### Development of predictor sets

The main goal of this study was to develop novel methods for classifying samples by radiation exposure type: “simple” exposures like homogeneous photon irradiation, *vs.* “complex” exposures like heterogeneous (*e.g.* partial-body) photon irradiation and/or neutron + photon mixed exposures. Therefore, in data set A we compared homogeneous and heterogeneous photon irradiation, and in data set B we compared photons only with neutron + photon mixtures.

Based on the distribution of micronuclei counts in each sample, we calculated several summary variables, described in Table 1, for evaluation as potential predictors of simple *vs.* complex exposure type. Heavily damaged cells are less likely to reach the binucleated state needed for micronuclei scoring, causing the total number of scored cells per sample to decrease with radiation dose. This phenomenon was the rationale for using the variable **LnSum**. The other variables listed in Table 1 were used based on our judgement of what metrics could act as reasonable potential predictors of exposure type and/or dose, combined with information about overdispersion of cytogenetic damage from complex exposure scenarios ^31, 38, 39^.

**Table 1.**
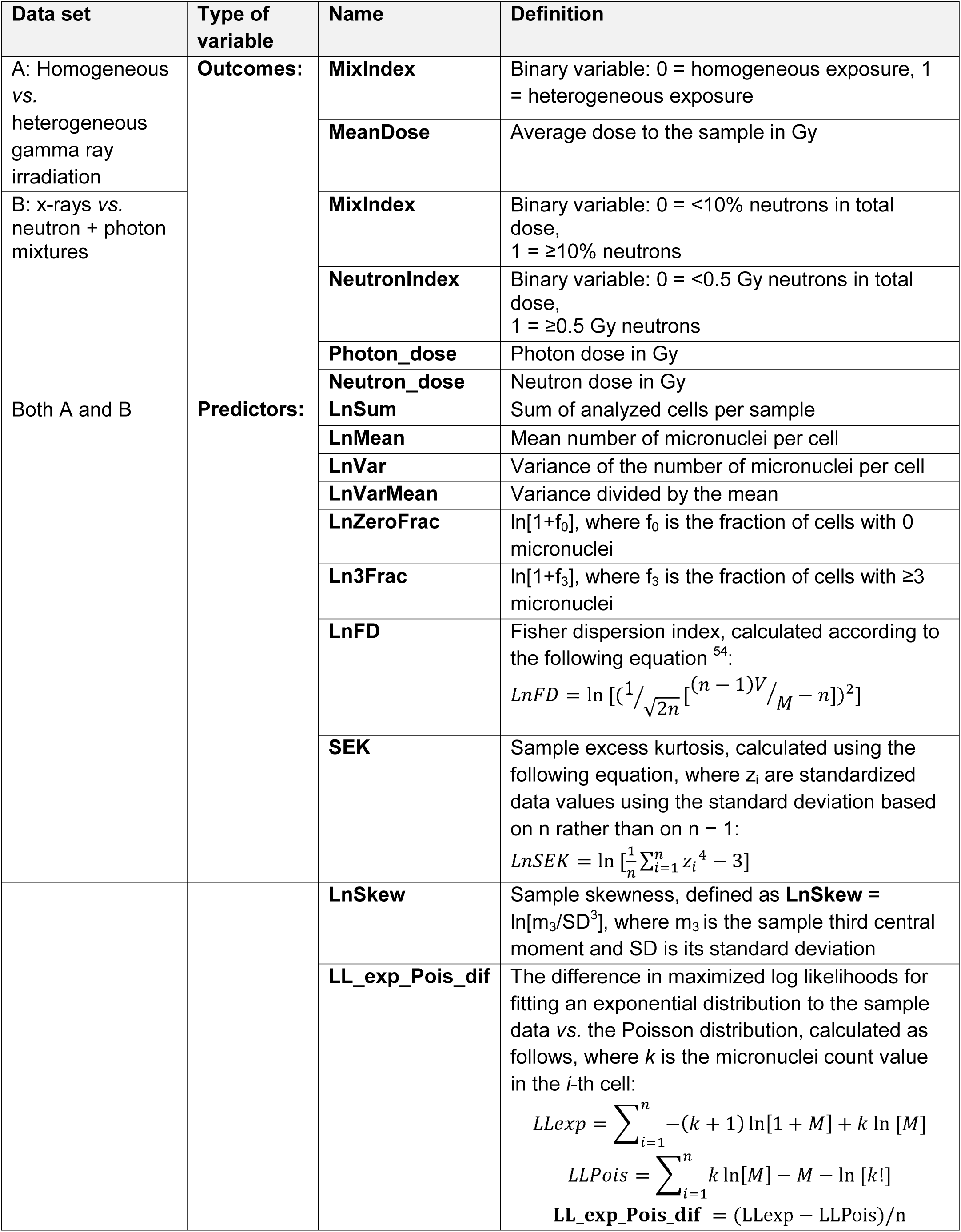
Descriptions of outcome (dependent) and predictor (independent) variables used in our analyses. The prefix “Ln” indicates natural logarithm. M is the mean, V is the variance, and n is the number of cells in the analyzed sample. The predictor variables were selected based on our judgement, combined with information about overdispersion of cytogenetic damage from complex exposure scenarios ^31, 38, 39^.

For data set A (homogeneous gamma irradiation of ex vivo human blood *vs.* 1:1 mixtures of irradiated and unirradiated blood) the outcome (independent) variables were called **MixIndex** and **MeanDose**. **MixIndex** was a binary variable, where 0 indicate homogeneous irradiation and 1 indicated a mixture of irradiated and unirradiated blood. **MeanDose** was the average gamma ray dose (in Gy), defined as the dose divided by 1+**MixIndex**. In other words, **MeanDose** for a sample of mixed blood was ½ of the dose received by the irradiated blood.

For data set B (ex vivo human blood irradiation with x-rays *vs.* neutron + photon mixtures) the outcome variables were called **Neutron_dose**, **Photon_dose**, **MixIndex**, and **NeutronIndex**. **Neutron_dose** and **Photon_dose** represent the dose contributions (in Gy) for each radiation type, respectively. The photon dose includes the gamma ray component of the neutron beam (∼18%) and the added x-ray dose. **MixIndex** in this data set was set to 1 if **Neutron_dose**/(**Neutron_dose**+ **Photon_dose**) ≥ 0.1, and set to 0 otherwise. **NeutronIndex** was set to 1 if **Neutron_dose** ≥ 0.5 Gy, and set to 0 otherwise. In other words, **MixIndex** = 1 indicated ≥10% neutron contribution to the total dose, and **NeutronIndex** = 1 indicated ≥0.5 Gy neutron dose. The cutoff values of 10% neutrons for **MixIndex** and 0.5 Gy for **NeutronIndex** were selected based on practical relevance and to create approximately balanced data classes (*i.e.* approximately equal numbers of samples above and below the cutoff). These outcome variables for both data sets are listed in Table 1. All parameter names starting with Ln are natural log transformed.

### Data analysis

We imported both data sets into *R* 3.5.1 software for analysis, and randomly split each of them into training and testing sets. Data set A was generated from a single experiment with a balanced design, with equal numbers of samples for homogeneous and heterogeneous radiation exposures. Consequently, we used the raw samples for analysis. In contrast, data set B was compiled from multiple experiments performed over several years, using a wide variety of photon and neutron doses. It contained 486 raw blood samples, where the total number of analyzed cells per sample varied greatly (from 33 to 3561) and the representation of different neutron + photon combinations was not equal. Consequently, we pooled (summed) all samples with the same combination of photon and neutron doses using the *aggregate* function in *R*. The raw and processed data sets are contained in the Supplementary_data_file1 online.

The training half of each data set was used for model fitting and selection, and the testing half was used to assess model performances. On the training data, we generated Spearman’s correlation coefficient matrices, including all predictors and outcome variables. To analyze all outcome variables simultaneously, using the same set of predictors, we employed the multivariate random forest (RF) machine learning approach (*MultivariateRandomForest R* package, https://cran.r-project.org/web/packages/MultivariateRandomForest/index.html) on each data set ^50^. The outcome variables were **MeanDose** and **MixIndex** for data set A, and **Neutron_dose**, **Photon_dose**, **MixIndex**, and **NeutronIndex** for data set B, as defined above. In data set B we also analyzed the “photon-equivalent dose”, defined as x-ray dose + RBE×neutron dose, where RBE is the neutron relative biological effectiveness. RBE was an adjustable parameter, and the analysis was performed using RF.

To focus in more detail on the main outcome variable of interest in both data sets, **MixIndex**, and to identify the strongest predictors of this variable, we also used the generalized boosted regression (GBM) algorithm ^45, 51^ (*gbm R* package, https://cran.r-project.org/web/packages/gbm/index.html) with a Bernoulli error distribution, and logistic regression (LR). The RF, GBM and LR methodologies and their implementation in our study are described in Supplementary Methods online.

## Results

### Analysis of data set A: homogeneous vs, non-homogeneous irradiation

#### Shape of micronucleus distribution

In this data set, partial-body exposures were mimicked by mixing gamma-irradiated and unirradiated blood samples with total-body exposures mimicked by standard ex-vivo irradiation. The goal of the analysis was to use metrics related to the shape of micronuclei per binucleated cell distributions to distinguish between homogeneous and mixed exposures. Differences in micronuclei/cell distributions between these exposure scenarios were apparent upon visual inspection of the pooled data (Fig. 2). For example, the distribution of micronuclei per cell for a 1:1 mixture of 4 Gy with 0 Gy irradiated blood was different from the distribution for blood irradiated with 2 Gy of pure gamma rays (Fig. 2), despite the fact that the mean micronuclei yields per binucleated cell were similar for these two scenarios (0.20 *vs.* 0.22, respectively).

**Figure 2.**
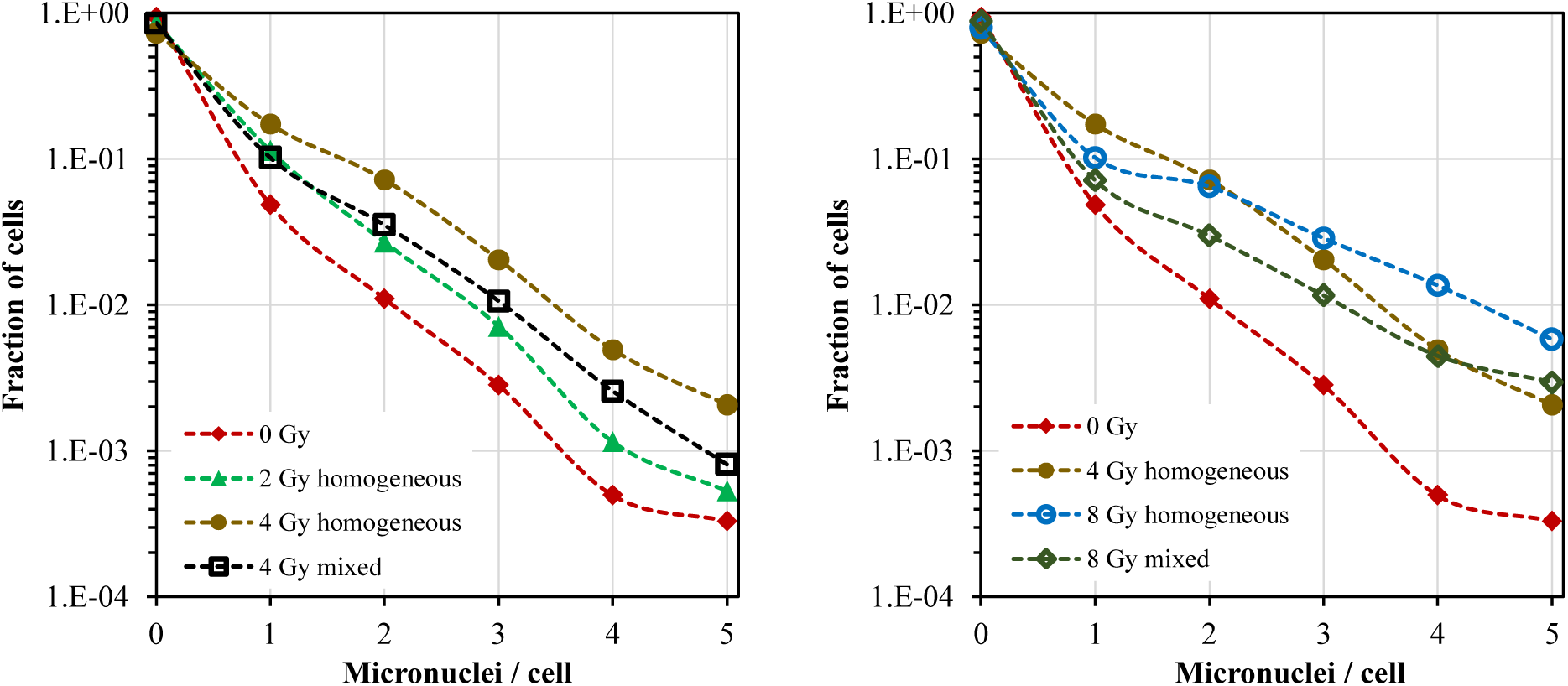
Distributions of micronuclei per binucleated cell data set A: blood samples ex vivo irradiated with 0, 2, 4 or 8 Gy of gamma rays (labeled “homogeneous”), or with 1:1 mixtures of 4 Gy with 0 Gy or 8 Gy with 0 Gy (labeled “mixed”). The differences between these distributions form the basis for our analysis aimed at discriminating between homogeneous and mixed exposures. Specifically, the data for 4 Gy mixed with 0 Gy are different from those for 2 Gy homogeneous (left panel), and the data for 8 Gy mixed with 0 Gy are different from those for 4 Gy homogeneous (right panel). Each curve was based on pooled analysis of a very large number of binucleated cells (from 8,417 to 21,056).

These differences were also reflected in the correlation matrix of predictors and outcomes (Fig. 3A). This matrix provides a convenient visualization of how all the analyzed variables are related to each other. As expected, the binary variable **MixIndex**, which indicated heterogeneous (mixed) *vs.* homogeneous exposure, was positively correlated with metrics of overdispersion: **LnVarMean**, **LnFD**, and **SEK** (Fig. 3A). In other words, overdispersed micronuclei/cell distributions with large “tails” were associated with heterogeneous exposures, whereas homogeneous irradiation was associated with lower variance/mean ratios and “tails”.

**Figure 3.**
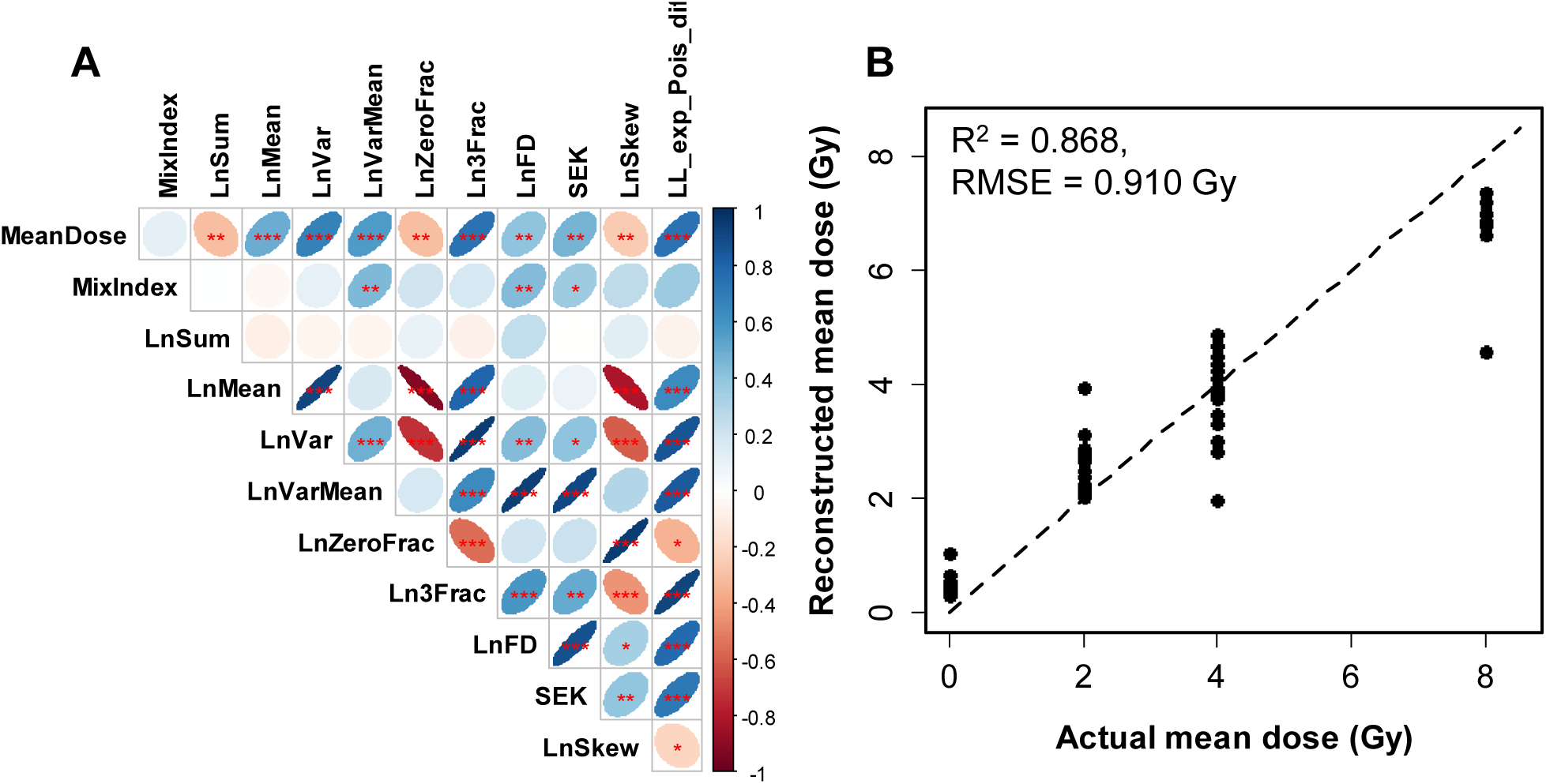
Analysis results summary for data set A: ex vivo human blood irradiated with homogeneous gamma ray doses *vs.* 1:1 mixtures of irradiated and unirradiated blood. **A.** Matrix of Spearman’s correlation coefficients (pairwise, without correction for multiple testing) between predictors and outcome variables. The meanings of all variables are provided in Table 1, and a color-coded correlation scale is provided on the right of the plot. Blue ellipses represent positive correlations, and red ones represent negative correlations. Darker color tones and narrower ellipses represent larger correlation coefficient magnitudes. Red star symbols indicate statistical significance levels: *** indicates p<0.001, ** indicates p<0.01, * indicates p<0.05, no stars indicates p>0.05. These p-values here are intended only for visualization: due to multiple comparisons, only 3 star significance levels are likely to indicate strong associations. Blank squares indicate correlation coefficients close to zero. **B.** Comparison of actual mean doses with reconstructed values by RF. Circles represent data points, and the line represents theoretically perfect 1:1 correlation.

The average dose received by each blood sample (**MeanDose**) was positively correlated with metrics for total damage, *e.g.* the mean micronuclei yield (**LnMean**) and the fraction of cells with ≥3 micronuclei (**Ln3Frac**), and negatively correlated with the sum of all analyzed cells (**LnSum**) and with the fraction of cells with zero micronuclei (**LnZeroFrac**) (Fig. 3A). In other words, the mean micronuclei yield, the total number of cells that made it to the binucleated stage, and the fraction of cells with no micronuclei were correlated with the average dose received by the blood sample.

#### Classification of partial body exposures

Multivariate machine learning analysis of data set A showed very good performance for reconstructing **MeanDose** and for detecting heterogeneous exposures (**MixIndex**) in a binary classification (Fig. 3B, Supplementary Table 1). Specifically, the area under the receiver operating characteristic curve (AUROC) for **MixIndex**, generated by RF analysis of on the testing data was 0.931 (range over 300 repeats was 0.903, 0.951), which falls into the “excellent” category for ROC curve metrics ^52^ (Supplementary Table 1). Univariate analyses using GBM and LR, which focused on reducing the predictor set and identifying the strongest predictors of **MixIndex**, as described in Supplementary Methods, performed in the “fair” to “good” range ^52^ (Supplementary Tables 1-2). The retained strongest predictors were **LL_exp_Pois_dif**, **LnVarMean**, and **LnFD** according to GB, and **LnFD** and **LL_exp_Pois_dif**×**SEK** according to LR. As mentioned above, these predictors indicate distribution shapes that are overdispersed relative to Poisson and are more similar to an exponential dependence, with a large “tail” at multiple micronuclei/cell. Their specific meanings are listed in Table 1 and in the Materials and Methods section.

### Analysis of data set B: photons vs. neutron + photon mixtures

This large data set consisted of ex vivo human blood samples exposed to x rays *vs.* neutron + photon mixtures in various proportions. The dependence of the mean micronucleus yield per binucleated cell on total radiation dose (photons + neutrons) and on the neutron contribution to this dose is shown graphically in Fig. 4. These data suggest that increasing the neutron contribution to the total dose notably increased the mean micronuclei yield, which is consistent with the high RBE of neutrons ^9–11^. It was also seen that, in mixed exposures, the yield of micronuclei is given by the sum of the yield of micronuclei we would expect from the separate photon and neutron irradiations – thus the two radiation types appear to be additive with respect to micronucleus yields.

**Figure 4.**
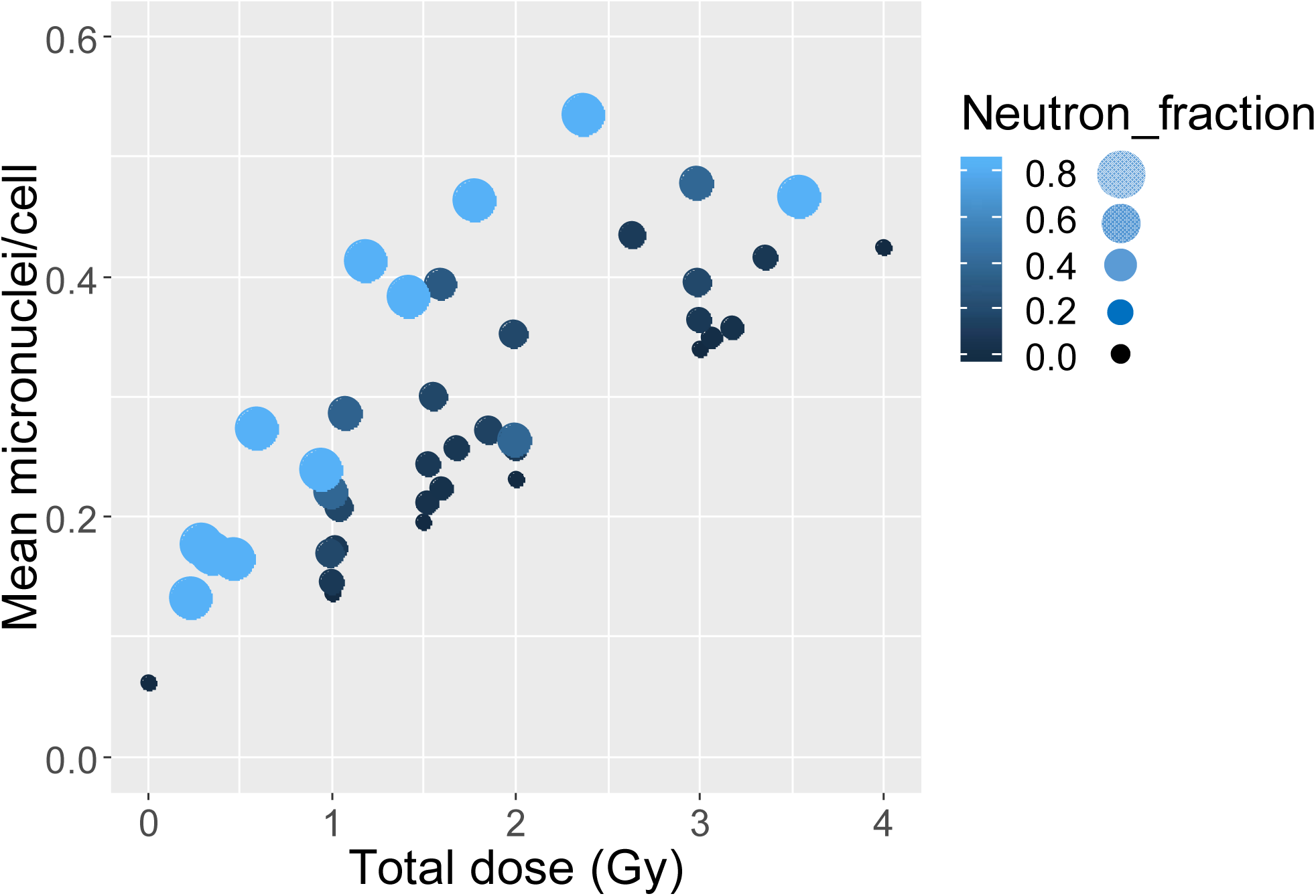
Dependence of mean micronuclei yield per binucleated cell on total radiation dose (photons + neutrons) and on the fraction of neutrons in this dose (Neutron_fraction). Larger and lighter colored circles represent a larger fraction of neutrons in the total dose.

#### Shape of micronucleus distribution

The presence of neutrons in the total dose also markedly alters the *shape* of the micronuclei per binucleated cell distributions. For example, Fig. 5 compares Poisson distribution fits to our micronuclei per cell data for 1.0 Gy of x-rays or 1.2 Gy of a neutron + photon beam (∼82% neutrons). The x-ray data in this example are clearly much more consistent with the Poisson distribution than the neutron beam data, which have a much larger “upper tail”, *i.e.* higher than Poisson-predicted probabilities of multiple micronuclei per cell.

**Figure 5.**
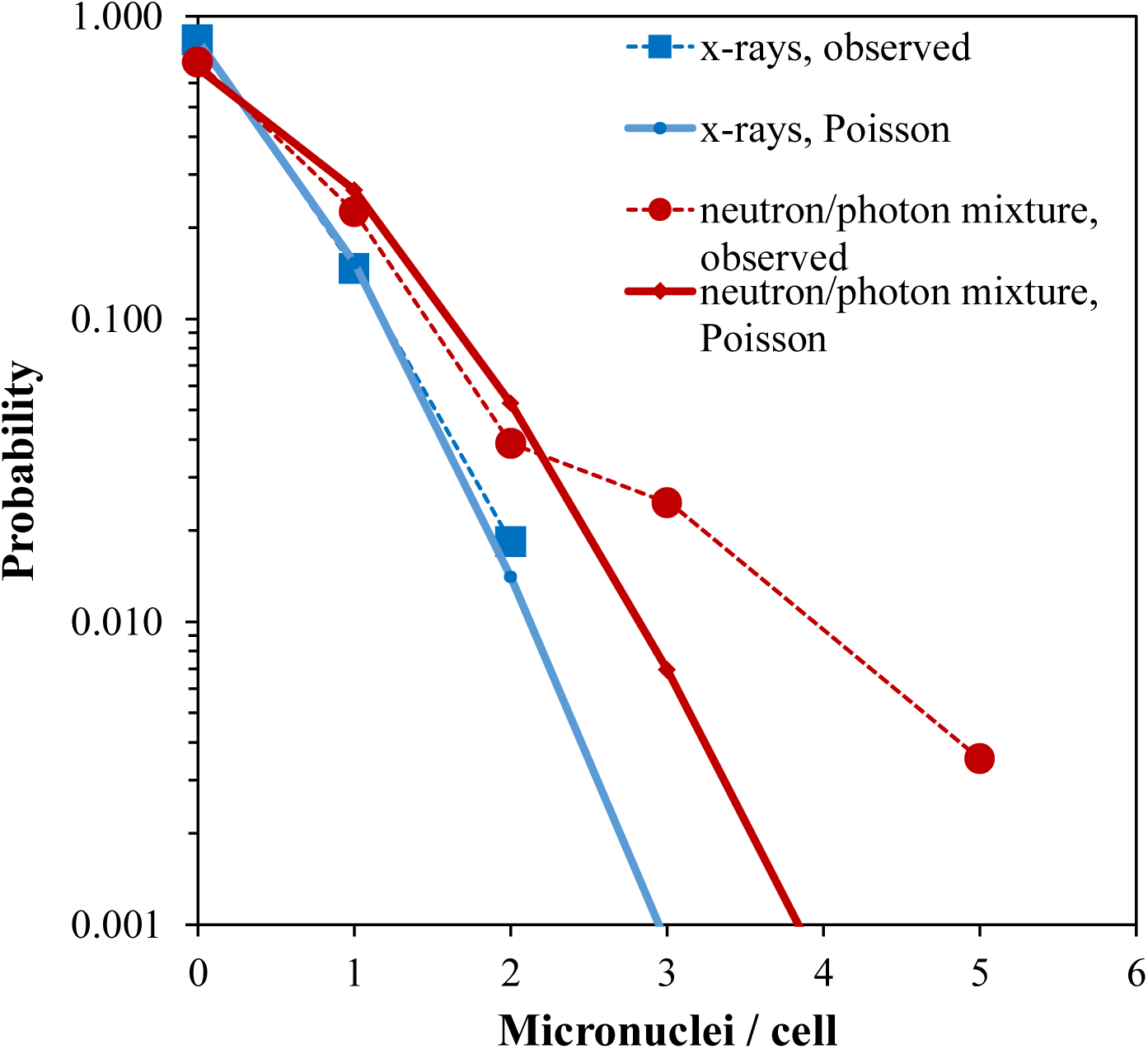
Comparison of Poisson distribution fits to micronuclei per binucleated cell data for 1.0 Gy x-rays *vs.* 1.2 Gy of a mixed neutron + photon beam that contains ∼82% neutrons. The probabilities of 3-5 micronuclei per cell in the mixed beam data are much larger, than those predicted by the best-fit Poisson distribution. No symbols are shown for micronuclei per cell values for which the observed counts were zero.

These effects of neutrons on the micronuclei/cell distribution are reflected in the correlation matrix of predictor and outcome variables are shown in Fig. 6A. Neutron dose was positively correlated with metrics for high damage yield (**LnMean**, **Ln3Frac**) and overdispersion (**LnVarMean**, **SEK**, **LL_exp_Pois_dif**), and negatively correlated with metrics for low damage yield (**LnSum**, **LnZeroFrac**) (Fig. 6A). Photon dose had the opposite correlation pattern regarding **LnVarMean**, **SEK** and **LL_exp_Pois_dif**, compared with neutron dose. These trends are intuitively explainable by the known overdispersion of neutron-induced damage compared with photon-induced damage ^38^.

**Figure 6.**
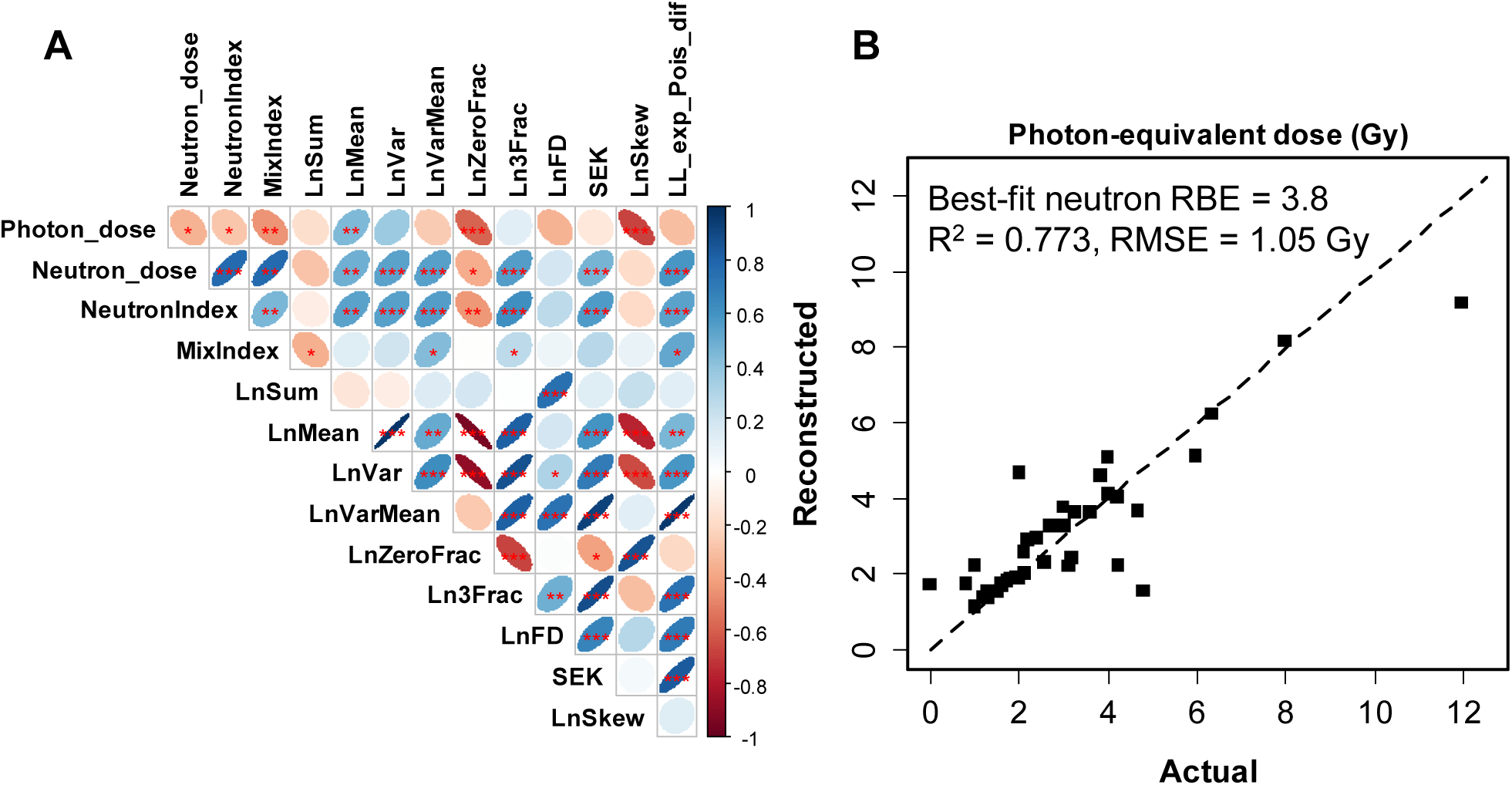
Analysis results summary for data set B: ex vivo human blood irradiated with x-rays *vs.* neutron + photon mixtures. **A.** Matrix of Spearman’s correlation coefficients (pairwise, without correction for multiple testing) between predictors and outcome variables. The meanings of all variables are provided in Table 1 and in the main text. The meanings of ellipse shapes and colors are the same as in Fig. 2, and a color-coded correlation scale is provided on the right of the plot. Blank squares indicate correlation coefficients close to zero. **B.** Comparison of actual photon-equivalent doses (defined as photon dose + RBE×neutron dose) with reconstructed values by RF. Circles represent data points, and the line represents theoretically perfect 1:1 correlation.

#### Classification of neutron exposures

The binary variable **NeutronIndex**, which indicated exposure to ≥0.5 Gy of neutrons, had essentially the same correlation patterns as neutron dose (Fig. 6A). The variable **MixIndex**, which indicated ≥10% of neutrons in the total dose, was most strongly positively correlated with two predictors: **LL_exp_Pois_dif** and **LnVarMean**, again suggesting that the overdispersion phenomenon is associated with neutron irradiation.

Multivariate RF analysis of data set B was quite good in reconstructing the photon-equivalent dose, defined as photon dose + RBE×neutron dose (Fig. 6B, Supplementary Table 1). The concordance between predictions and actual values was particularly close in the dose region around 2 Gy, which is important for triage decision-making (Fig. 6B). The best-fit neutron RBE value was 3.8, very similar to the previously published value of 4 for micronuclei following irradiation at CINF ^9^.

Notably, multivariate RF was very good at detecting a neutron fraction ≥10% (**MixIndex**) and neutron doses ≥0.5 Gy (**NeutronIndex**) in binary classifications (Fig. 7A-B). The AUROC values for **MixIndex** and **NeutronIndex** were 0.916 (uncertainty range 0.893 to 0.943 over 300 RF repeats) and 0.848 (0.815 to 0.879), respectively (Supplementary Table 1). These values fall into the good to excellent range for ROC curve metrics ^52^. Targeted analyses using GBM and LR (described in Supplementary Methods) performed as well as RF in predicting **MixIndex**, with AUROC of 0.922 (0.878, 0.961) and 0.911 (0.819, 1.0), respectively (Supplementary Tables 1, 3). These techniques used fewer predictors: **LnVarMean**, **LL_exp_Pois_dif**, **LnSum**, **SEK**, **Ln3Frac**, and **LnZeroFrac** for GB, and **LnSum**, **LL_exp_Pois_dif**×**Ln3Frac**, and **LL_exp_Pois_dif**×**LnSum** for LR. Therefore, accurate predictions of **MixIndex** were generated using predictor groups that were indicative of overdispersion (*e.g.* **LnVarMean** and **LL_exp_Pois_dif**) and total damage yields (*e.g.* **LnSum**, **Ln3Frac**, and **LnZeroFrac**).

**Figure 7.**
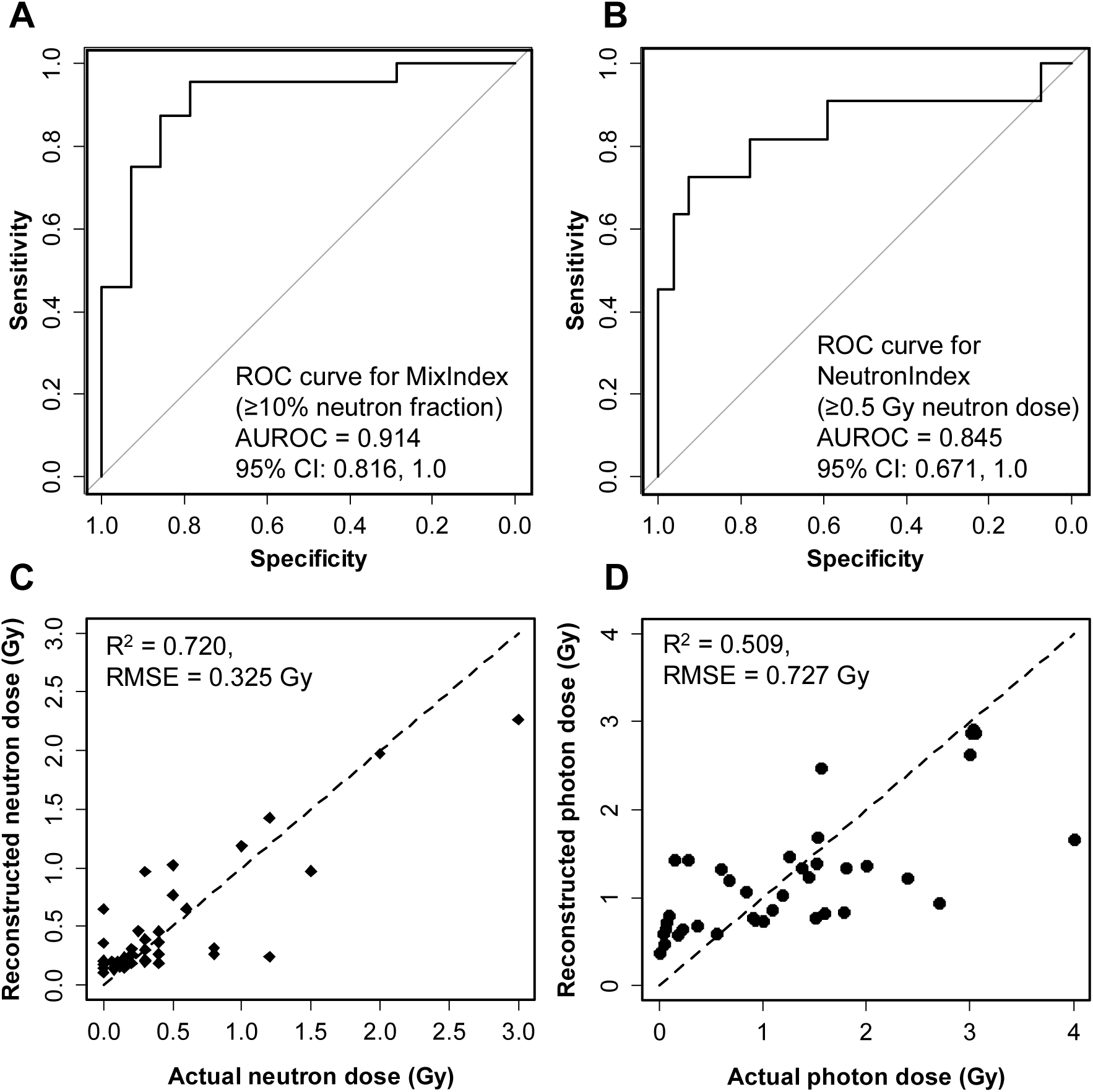
RF performance for data set B: ex vivo human blood irradiated with x rays *vs.* neutron + photon mixtures. **A.** ROC curve for discriminating between exposures with ≥10% neutron fraction *vs.* those with <10% neutrons. **B.** ROC curve for discriminating between exposures with ≥0.5 Gy neutron dose *vs.* those with <0.5 Gy neutrons. **C-D.** Comparisons of actual and reconstructed neutron and photon doses, respectively. Circles represent data points, and the lines represents theoretically perfect 1:1 correlation.

Quantitative reconstructions of the neutron and photon dose components (**Neutron_dose** and **Photon_dose**, respectively) were weaker, compared with the binary classifications. Neutron dose reconstructions were decent (Fig. 7C, Supplementary Table 1), and photon dose reconstructions were relatively poor (Fig. 7D, Supplementary Table 1). These results may indicate that the selected predictor set, which was focused on micronuclei/cell distribution shapes, is a sensitive qualitative indicator of complex exposure scenarios, but is less sensitive for quantifying the details of these scenarios.

## Discussion

The CBMN assay is one of the simplest cytogenetic biodosimetry assays to perform and score. It is therefore also the easiest to automate ^53^. However, the conventional CBMN assay is geared towards uniform photon exposures and provides only the photon-equivalent total body dose, which may not be the most useful parameter in scenarios involving mixed neutron + photon or partial body exposures. To address these issues, there are many advanced techniques for dose reconstruction for complex exposure scenarios, which are generally based on fitting parametric linear or linear quadratic dose response functions with selected error distributions (*e.g.* Zero-Inflated Poisson or Negative Binomial) ^31, 40, 42, 43^. Here we extended the analysis of micronuclei per cell distribution shapes in a different direction: we used various summary metrics like index of dispersion, skewness and kurtosis as potential predictors of complex exposure scenarios, and imported these predictors into machine learning or parametric regression methods.

The conceptual basis for our approach is that micronuclei per binucleated cell distributions from complex exposures have different shapes (*e.g.* “tails”), compared with distributions from simple exposures, even when the mean micronucleus yields are the same for both scenarios. These differences in distribution shapes translated into differences in variables like index of dispersion, kurtosis and skewness (Table 1), which were used to generate predictors imported into machine learning and parametric modeling approaches. To our knowledge, this approach is new and was not used previously in radiation biodosimetry.

Our results suggest that 1:1 mixtures of irradiated and unirradiated blood can be quite accurately discriminated from homogeneous irradiations (AUROC > 0.9 on testing data, Supplementary Table 1). Ongoing work is focusing on determination of the minimal shielded percentage that can be reliably detected.

Using the same approaches, we also obtained encouraging results in discrimination of mixed exposures to photons and neutrons from pure photon exposures, *e.g.* by detecting ≥10% neutron fractions or ≥0.5 Gy of neutrons in the total dose (AUROC > 0.9 for the first scenario and > 0.8 for the second, Fig. 7A-B, Supplementary Table 1). Of note, the dose reconstructions performed using this method estimated the measured RBE rather well (3.8 in this work *vs.* 4 in reference ^9^). Ongoing work focuses on obtaining more precise reconstructions of the neutron fractions and photon doses.

Therefore, although the two scenarios (partial body and neutron exposures) differ in experimental design and radiation doses and types, the general concept of using micronucleus distribution shape metrics as indicators of complex *vs.* simple exposure scenarios was applicable in both situations. At this stage, our results of course represent only a proof of principle because ex vivo blood irradiation is an “idealized” model system for partial-body and neutron + photon mixed exposures. Much more complexity is expected for realistic in vivo scenarios because various organs, which are (or are not) irradiated in the ex vivo situation, can contribute to the in vivo responses. Furthermore, a realistic exposure may include both neutron and partial body photon exposures. These type of scenarios were not investigated in this work but will be the focus of future studies. The accuracy of applying the approaches proposed here under realistic mass-casualty conditions can probably be increased by integrating micronuclei assays with other types of radiation biomarkers (*e.g.* dicentric chromosomes, gene expression levels, blood cell counts). Several biomarkers, combined into one framework, are likely to provide more detailed and useful information than a single assay alone.

### Conclusions

We demonstrate a proof of principle that measurements of the distributions of micronuclei per binucleated cell, analyzed by a novel implementation of machine learning and parametric regression methods, contain enough information to detect complex exposure scenarios involving partial-body shielding or densely ionizing radiations. The ability to perform such detection reliably in a high throughput manner would be extremely useful in radiation-related mass casualty situations such as IND detonations because partial-body and/or neutron exposures can have very different clinical outcomes, compared with homogeneous photon irradiation.

## Supporting information

Data sets

## Author Contributions

IS, HCT, JRP, LC, MPC, MHD, AH, AB, MT, GG, and DJB conceived the study and prepared the manuscript. IS carried out the data analysis. HCT, JRP, LC, MPC, MHD, AH, AB, MT, and GG performed the experimental studies. All authors contributed to editing the manuscript.

## Additional Information

**Supplementary information** accompanies this paper

**Competing interests:** The authors declare no competing interests.

**Funding:** This work was supported by the National Institute of Allergy and Infectious Diseases grant 5U19AI067773-14.

## List of supplementary files

**Supplementary data file 1:** Data sets used for analysis

**Supplementary tables:** Supplementary tables 1-3.

**Supplementary methods:** Supplementary methods.

