## Supplementary tables for "New High Throughput Approaches to Detect Partial-body and Neutron Exposures on an Individual Basis"

**Supplementary Table 1.** **Summary performance metrics for multivariate random forest (RF), generalized boosted regression (GBM), and logistic regression (LR) analyses of different outcome variables in both data sets.** AUROC = area under the ROC curve. RMSE = root mean squared error (here in units of Gy). The uncertainty range represents DeLong 95% CIs ^1^ for logistic regression, and range over 300 repeats with different initial random number seeds for RF and GBM.

| **Data set** | **Outcome variable** | **Data analysis method** | **Perfor-mance metric** | **Mean** | **Uncertainty range** | |
| --- | --- | --- | --- | --- | --- | --- |
| A: Homogeneous *vs.* heterogeneous gamma ray irradiation | MixIndex  (0 = homogeneous, 1 = heterogeneous) | RF | AUROC | 0.931 | 0.903 | 0.951 |
|  |  | GBM |  | 0.846 | 0.805 | 0.886 |
|  |  | LR |  | 0.754 | 0.611 | 0.896 |
|  | MeanDose  (average dose to the sample in Gy) | RF | R^2^ | 0.869 | 0.833 | 0.904 |
|  |  |  | RMSE | 0.924 | 0.822 | 1.021 |
| B: x-rays *vs.* neutron/photon mixtures | MixIndex  (0 = <10% neutrons,  1 = ≥10% neutrons) | RF | AUROC | 0.916 | 0.893 | 0.943 |
|  |  | GBM |  | 0.922 | 0.872 | 0.961 |
|  |  | LR |  | 0.911 | 0.819 | 1.000 |
|  | NeutronIndex  (0 = <0.5 Gy neutrons,  1 = ≥0.5 Gy neutrons) | RF | AUROC | 0.848 | 0.815 | 0.879 |
|  | Photon_dose  (photon dose in Gy) | RF | R^2^ | 0.513 | 0.478 | 0.557 |
|  |  |  | RMSE | 0.724 | 0.694 | 0.749 |
|  | Neutron_dose  (neutron dose in Gy) | RF | R^2^ | 0.706 | 0.666 | 0.736 |
|  |  |  | RMSE | 0.333 | 0.316 | 0.354 |

**Supplementary Table 2.** **Summary of GBM performance results (over 300 repeats) in detecting heterogeneous *vs.* homogeneous exposures (MixIndex) in data set A.** The error index represents misclassifications, calculated using the mean GBM predictions for the given sample and an adjustable cutoff value of 0.55, which was selected to minimize the overall error rate. SD represents standard deviation. The misclassification rate (average error index) for homogeneously exposed samples (those with MixIndex = 0) was 25.0%, and for heterogeneously exposed samples (those with MixIndex = 1) it was 15.0%.

| **Mean dose (Gy)** | **MixIndex** | **GBM predictions for MixIndex** | | **Error index** |
| --- | --- | --- | --- | --- |
|  |  | **Mean** | **SD** |  |
| 0 | 0 | 0.101 | 0.051 | 0 |
| 0 | 0 | 0.072 | 0.034 | 0 |
| 2 | 0 | 0.097 | 0.039 | 0 |
| 2 | 0 | 0.930 | 0.033 | 1 |
| 2 | 0 | 0.503 | 0.082 | 0 |
| 4 | 0 | 0.522 | 0.094 | 0 |
| 4 | 0 | 0.737 | 0.071 | 1 |
| 4 | 0 | 0.543 | 0.083 | 0 |
| 4 | 0 | 0.487 | 0.083 | 0 |
| 8 | 0 | 0.672 | 0.070 | 1 |
| 8 | 0 | 0.173 | 0.075 | 0 |
| 8 | 0 | 0.166 | 0.075 | 0 |
| 0 | 0 | 0.120 | 0.053 | 0 |
| 0 | 0 | 0.111 | 0.052 | 0 |
| 0 | 0 | 0.858 | 0.053 | 1 |
| 0 | 0 | 0.136 | 0.057 | 0 |
| 0 | 0 | 0.148 | 0.056 | 0 |
| 2 | 0 | 0.552 | 0.091 | 1 |
| 2 | 0 | 0.708 | 0.068 | 1 |
| 4 | 0 | 0.288 | 0.079 | 0 |
| 4 | 0 | 0.470 | 0.088 | 0 |
| 4 | 0 | 0.701 | 0.070 | 1 |
| 4 | 0 | 0.533 | 0.080 | 0 |
| 8 | 0 | 0.353 | 0.089 | 0 |
| 8 | 0 | 0.346 | 0.093 | 0 |
| 8 | 0 | 0.375 | 0.081 | 0 |
| 8 | 0 | 0.499 | 0.070 | 0 |
| 8 | 0 | 0.385 | 0.074 | 0 |
| 2 | 1 | 0.933 | 0.031 | 0 |
| 2 | 1 | 0.944 | 0.027 | 0 |
| 2 | 1 | 0.893 | 0.044 | 0 |
| 4 | 1 | 0.257 | 0.079 | 1 |
| 4 | 1 | 0.957 | 0.024 | 0 |
| 4 | 1 | 0.924 | 0.035 | 0 |
| 2 | 1 | 0.622 | 0.072 | 0 |
| 2 | 1 | 0.871 | 0.048 | 0 |
| 2 | 1 | 0.884 | 0.046 | 0 |
| 4 | 1 | 0.898 | 0.051 | 0 |
| 2 | 1 | 0.737 | 0.076 | 0 |
| 2 | 1 | 0.559 | 0.075 | 0 |
| 4 | 1 | 0.847 | 0.055 | 0 |
| 2 | 1 | 0.899 | 0.040 | 0 |
| 2 | 1 | 0.224 | 0.073 | 1 |
| 4 | 1 | 0.853 | 0.044 | 0 |
| 4 | 1 | 0.513 | 0.073 | 1 |
| 4 | 1 | 0.827 | 0.055 | 0 |
| 4 | 1 | 0.600 | 0.070 | 0 |
| 4 | 1 | 0.932 | 0.028 | 0 |

**Supplementary Table 3.** **Summary of GBM performance results (over 300 repeats) in detecting ≥10% neutron fraction of the total dose (MixIndex) in data set B.** The error index represents misclassifications, calculated using the mean GBM predictions for the given sample and an adjustable cutoff value of 0.6, which was selected to minimize the overall error rate. SD represents standard deviation. The misclassification rate (average error index) for homogeneously exposed samples (those with MixIndex = 0) was 21.4%, and for heterogeneously exposed samples (those with MixIndex = 1) it was 4.2%.

| **Neutron dose (Gy)** | **MixIndex** | **GBM predictions for MixIndex** | | **Error index** |
| --- | --- | --- | --- | --- |
|  |  | **Mean** | **SD** |  |
| 0.000 | 0 | 0.782 | 0.090 | 1 |
| 0.000 | 0 | 0.267 | 0.096 | 0 |
| 0.000 | 0 | 0.793 | 0.093 | 1 |
| 0.000 | 0 | 0.214 | 0.099 | 0 |
| 0.000 | 0 | 0.084 | 0.066 | 0 |
| 0.000 | 0 | 0.683 | 0.122 | 1 |
| 0.060 | 0 | 0.342 | 0.116 | 0 |
| 0.075 | 0 | 0.549 | 0.134 | 0 |
| 0.075 | 0 | 0.294 | 0.095 | 0 |
| 0.100 | 0 | 0.411 | 0.126 | 0 |
| 0.150 | 0 | 0.175 | 0.104 | 0 |
| 0.150 | 0 | 0.429 | 0.117 | 0 |
| 0.150 | 0 | 0.480 | 0.132 | 0 |
| 0.300 | 0 | 0.530 | 0.140 | 0 |
| 0.100 | 1 | 0.914 | 0.051 | 0 |
| 0.200 | 1 | 0.950 | 0.037 | 0 |
| 0.200 | 1 | 0.666 | 0.116 | 0 |
| 0.200 | 1 | 0.681 | 0.110 | 0 |
| 0.250 | 1 | 0.950 | 0.037 | 0 |
| 0.300 | 1 | 0.648 | 0.114 | 0 |
| 0.300 | 1 | 0.955 | 0.033 | 0 |
| 0.300 | 1 | 0.238 | 0.103 | 1 |
| 0.300 | 1 | 0.974 | 0.025 | 0 |
| 0.400 | 1 | 0.769 | 0.079 | 0 |
| 0.400 | 1 | 0.666 | 0.117 | 0 |
| 0.400 | 1 | 0.952 | 0.039 | 0 |
| 0.400 | 1 | 0.947 | 0.040 | 0 |
| 0.500 | 1 | 0.974 | 0.024 | 0 |
| 0.500 | 1 | 0.938 | 0.043 | 0 |
| 0.600 | 1 | 0.973 | 0.025 | 0 |
| 0.800 | 1 | 0.786 | 0.097 | 0 |
| 0.800 | 1 | 0.941 | 0.040 | 0 |
| 1.000 | 1 | 0.936 | 0.042 | 0 |
| 1.200 | 1 | 0.931 | 0.044 | 0 |
| 1.200 | 1 | 0.973 | 0.025 | 0 |
| 1.500 | 1 | 0.971 | 0.027 | 0 |
| 2.000 | 1 | 0.965 | 0.032 | 0 |
| 3.000 | 1 | 0.937 | 0.043 | 0 |
