## Supplementary methods for "New High Throughput Approaches to Detect Partial-body and Neutron Exposures on an Individual Basis"

Random forests (RF) and generalized boosted regression (GBM) are powerful machine learning techniques that make predictions based on ensembles of decision trees. RF uses decision trees as base models, and employs “bagging” and tree de-correlation approaches to improve performance. The bagging (bootstrapping and aggregation) procedure involves generating bootstrapped samples and using a random subsample of the features for each fitted decision tree. Decision trees have some very useful properties for analyzing noisy data sets with different predictor types and a large ratio of predictors to observations. They are not sensitive to outliers and to the presence of many weak or irrelevant predictors. They are also unaffected by monotonic (*e.g.* logarithmic) transformations of the data. GBM also uses decision trees, but the trees are averaged by boosting rather than bagging. Boosting involves iterative fitting of trees: the data are reweighted so that the next trees focus more strongly on those data points on which previous trees performed the worst. GBM readily accommodates different types of error distributions, *e.g.* Gaussian for continuous data and Bernoulli for binary data. The historically noted tendency of tree-based machine learning algorithms to overfit the data was alleviated by using independent testing data sets, repeated cross validation on training data sets, and by preventing the terminal nodes in any tree from containing less than 5-10% of the sample size. The tuning parameters for RF (number of trees in the forest, number of randomly selected features considered for a split in each regression tree node, and minimum number of samples in the leaf node) and for GBM (total number of trees to fit, highest level of variable interactions, minimum number of observations in the terminal nodes, and the shrinkage or learning rate) were optimized using the *caret* *R* package (https://cran.r-project.org/web/packages/caret/caret.pdf) with repeated 10-fold cross validation, or by a customized *R* code that looped over multiple values of each parameter and evaluated model performance.

Once optimal tuning parameter values were found, RF and GBM analyses were repeated 300 times for each data set using different initial random number seeds to assess the robustness of predictions. For binary outcome variables like MixIndex (both data sets) and NeutronIndex (data set B), receiver operating characteristic (ROC) curves were constructed, and the model performance metric was area under the ROC curve (AUROC), estimated using the *pROC* *R* package (https://cran.r-project.org/web/packages/pROC/pROC.pdf). For continuous outcome variables (MeanDose in data set A, Photon_dose and Neutron_dose in data set B) the performance metrics were the coefficient of determination (R^2^) and root mean squared error (RMSE) for comparing real with predicted doses. The last of 300 RF repeats was used as a typical example to generate graphical representations of how the models fitted the data.

As expected, in the various methods implemented here some predictor variables were more influential on the predictions, than others. To identify the most important predictors, we examined relative influence scores from GBM analyses and sequentially removed those predictors that achieved the lowest average scores across GBM repeats. This process of manual removal of weak predictors was continued until removing one more predictor dramatically lowered the average AUROC value across GBM repeats.

We also performed systematic feature selection in the context of parametric logistic regression (LR) on both data sets, with MixIndex as the outcome variable. Some of the predictor variables were, as expected, strongly correlated to each other, potentially resulting in multicollinearity. We sought to alleviate this problem by stepwise removal of predictor variables with high variance inflation factors (VIFs) until a smaller predictor set with lower VIFs (<5) was generated for each data set. Using the reduced predictor set, we performed multimodel inference (MMI) with the *glmulti* *R* package (https://cran.r-project.org/web/packages/glmulti/glmulti.pdf). This approach evaluates all possible main effects and pairwise interactions of the predictors and assigns relative importance scores to each of these predictor combinations based on the Akaike information criterion with sample size correction (AICc). Uncertainties (95% confidence intervals, CIs) with correction for model selection uncertainty were generated by this method. Those predictors that had 95% CIs that do not overlap zero and/or had the highest importance scores were considered as the strongest ones, and they were retained in a preferred logistic regression model. The preferred model was tested on the testing data by calculating AUROC.
